# A Comprehensive Survey of Mutations in Oesophageal Carcinoma Reveals Recurrent Neoantigens as Potential Immunotherapy Targets

**DOI:** 10.1101/2020.03.28.013201

**Authors:** Chao Chen, Songming Liu, Heng Xiong, Xi Zhang, Bo Li

## Abstract

This study was aimed to investigate the mutations in Esophageal Carcinoma (EC) for recurrent neoantigen identification. A total of 733 samples with whole exome sequencing (WES) mutation data and 1153 samples with target region sequencing data were obtained from 7 published studies and GENIE database. Common HLA-I and HLA-II genotypes in both TCGA cohort and Chinese were used to predict the probability of ‘public’ neoantigens in the dataset. Based on the integrated data, we not only obtained the most comprehensive EC mutation landscape so far, but also found 253 mutation sites which could be identified in at least 3 or more patients, including, *TP53* p.R248Q, *PIK3CA* p.E545K, *PIK3CA* p.E542K, *KRAS* p.G12D, *PIK3CA* p.H1047R and *TP53* p.C83F. These mutations can be recognized by multiple common HLA molecules (HLA-A11:01, HLA-B57:01, HLA-A03:01, DRB1-0301, DRB1-1202, et al.) in Chinese and TCGA cohort as potential public neoantigens. Overall, our analysis provides some potential targets for EC immunotherapy.

## 1. Introduction

Esophageal cancer (EC) is one of the most common malignant tumors in the world, especially in south China [1]. Chemotherapy, anti-HER2 and immune checkpoint inhibitors (ICIs) therapy are the most popular treatment strategies for EC, but for advanced cancer, the effect of these treatment methods is still limited, and the 5-year survival rate of metastatic EC is less than 5%[2]. Therefore, it is necessary to further explore other treatment methods.

EC is mainly divided into two subtypes, esophageal squamous cell carcinoma (ESCC) and esophageal adenocarcinoma (EAC). ESCC is the main subtype of EC, accounting for about 90% of all esophageal cancers [3]. With the development of the next-generation sequencing (NGS) technology, the genomics research of ESCC and EAC has been widely carried out, and the molecular characteristics and some key oncogenes in their subtypes have been found[3–9]. The main purpose of these studies is to find out the mutation characteristics and driver genes of EC, less involved neoantigen-based immunotherapy, and because of the limited number of samples, it is difficult to find the common high-frequency neoantigens among EC patients. In recent years, immunotherapy based on tumor-specific neoantigens has made remarkable progress in solid tumor therapy. It has been reported that tumor specific T cells has achieved therapeutic effect in melanoma[10, 11], breast cancer[12] and malignant glioma[13]. However, as far as we know, there is few reports on the application of T-cell therapy based on tumor-specific neoantigens in EC.

In this study, by integrating the previous 6 WES study data and panel mutation data of 1153 EC patients in GENIE database, the mutation data of 1886 EC patients were finally obtained. Furthermore, we screened the high frequency mutation sites and used them to predict neoantigens. Finally, we got some common neoantigen targets among EC patients, which will be verified in subsequent immunogenicity or functional experiments.

## 2. Materials and Methods

### 2.1. Genomic Data of EC

This study was approved by the Institutional Review Board on Bioethics and Biosafety of BGI group. All somatic mutations, including single nucleotide variants (SNVs) and short insertion/deletion (indels), were downloaded from the latest publications (Table 1 and Supplementary Table S1), which represent six geographically diverse study groups involving 733 EC patients. Since all data used in this study were from public databases with informed consent from participants in the original genome study, no additional informed consent was required.

**Table 1:**
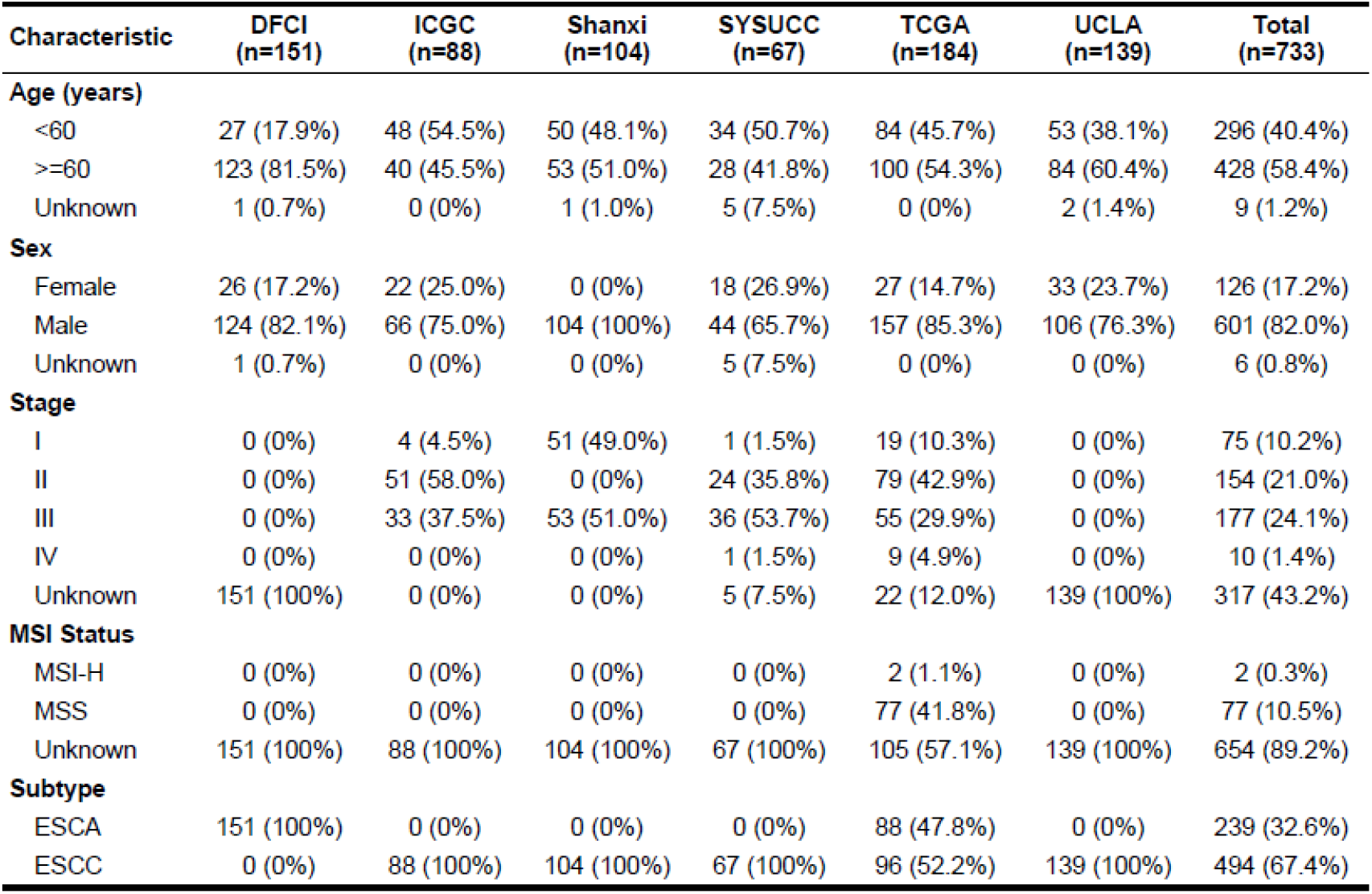
Summary of clinical information including patients from six studies and 733 EC samples.

### 2.2. Pipeline for Neoantigen Prediction

For neoantigen prediction, a total of 43 HLA-I genotypes and 8 HLA-II genotypes were selected for neoantigen affinity and presentation probability assessment as previously reported (Supplementary Table S2)[14]. Mutations, which include 214 non-silent somatic SNVs and 39 indels, could be identified in at least 3 patients. All these mutations were then used for neoantigen prediction. For HLA-I alleles, neoantigens were predicted by NetMHC[15], NetMHCpan[16], PickPocket[17], PSSMHCpan[18], SMM[19] and EPIC[20]. According to our previous research[21], neoantigen peptides need to meet the following four criteria: (1) Between 8-11 mers length; (2) Affinity IC50 < 500nM on at least two tools; (3) Mutant (MT) peptides affinity of score lower than the wild type (WT); (4) The rendering score in EPIC is greater than 0.5. For HLA-II alleles, neoantigens were predicted by netMHCIIpan[16], peptides need to meet the following criteria: (1) Between 13-25 mers length; (2) Affinity IC50 < 50nM or the Strong Binders: rank<0.5%.

### 2.3. Statistical Analysis

The statistical analysis was done in R-studio and the mutation analysis and drawing were done with the maftools package[22]. If no special specification were given, P < 0.05 was considered significant.

## 3. Results and Discussion

### 3.1. The Integrated Mutation Landscape of EC Patients

The mutation status of all samples is shown in **Figure 1 and Figure S1**. In general, missense mutation is the most main type of mutations. At the base substitution level, C>T is the dominant mutant form, followed by C>A and C>G (**Figure S2**), which is consistent with TCGA and previous reports[4, 7]. The median number of mutations in each sample was 81, among which *TP53, TTN, MUC16, SYNE1, CSMD3, PCLO*, and *LRP1B* were the most frequently mutated genes. In all 733 samples, 239 samples of esophageal adenocarcinoma (EAC) and 494 samples of esophageal squamous carcinoma (ESCC), the overall mutation burden of EAC was significantly higher than ESCC, the median mutation number of EAC and ESCC were 117 and 58, respectively (**Figure S3-S4**).

**Figure 1.**
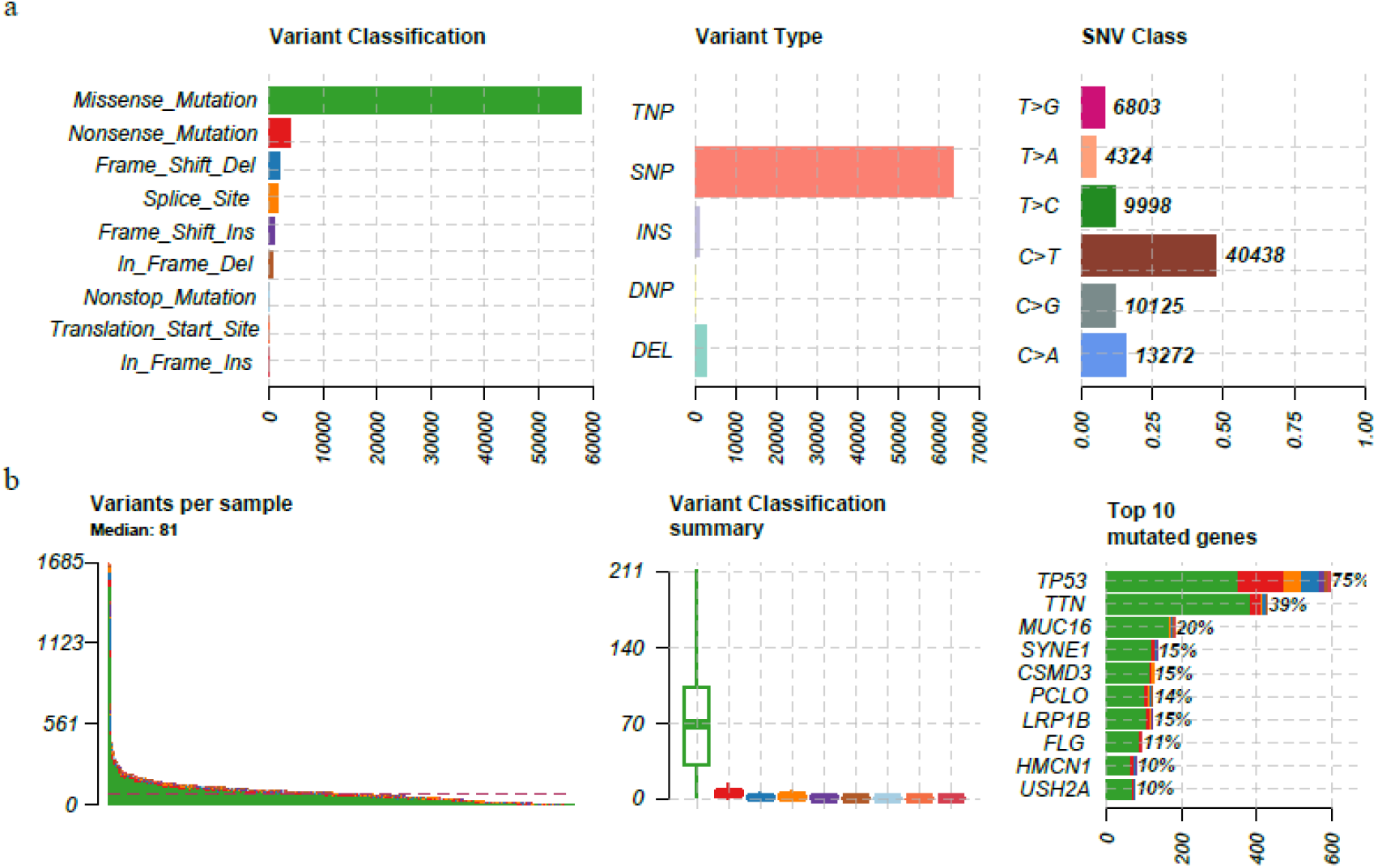
The mutation landscape in EC cohort. (a) from left to right, counts of each variant classification, counts of each variant type, counts of each SNV class. (b) from left to right, variants number per sample, variant classification and top 10 significantly mutated genes.

Interestingly, through the integration analysis of mutation data of 733 WES samples and 1153 panel samples in GENIE database, we found that mutations of multiple genes, such as *MUC16, FAT3 and CDKN2A, PIK3CA* and *TP53*, were mutually exclusive (Fisher’s exact test, P<0.05, **Figure S5**). This different mutation pattern may suggest that the carcinogenic mechanisms are different in EC patients carrying mutations in these genes. It’s worth noting that the mutual exclusion of *PIK3CA* and *TP53* also exists in colorectal cancer and gastric cancer[14, 23, 24], indicating that this may be a common feature of digestive system cancer.

There are many hot spot mutations in EC samples. Of these, 253 recurrent mutations could be identified in at least 3 patients, with 214 SNVs and 39 indels, respectively. Previous studies have shown that common neoantigens in cancers could be used as potential immunotherapy targets[21, 23]. Therefore, in order to find out whether there are common neoantigens in EC populations, we used these mutations in downstream analyses to predict tumour-specific neoantigens.

### 3.2. Neoantigen Profiling of EC Patients

Due to the difference frequency of HLA-I and HLA-II genes in different populations, in order to search for ‘public’ neoantigens in EC populations, we selected high-frequency HLA alleles in Chinese (HLA frequency >5% in Han Chinese [25]) and high-frequency HLA in western people (HLA frequency >5% in TCGA [26]) for neoantigen analysis. Finally, a total of 43 HLA-I and 8 HLA-II alleles were used for neoantigen prediction (**Table S2**).

We detected 473 SNV derived neoantigens and 111 INDEL derived neoantigens (**Table S3-4**). Each SNV usually produces 1-2 high-affinity peptides, while each indel can produce multiple high-affinity peptides. The top fifteen high frequency mutation sites and neoantigen peptides in SNV and indel are shown in **Table 2** and **Table 3**, respectively. In terms of SNV, mutations of *TP53, KRAS, PIK3CA* and *CDKN2A* can produce neoantigens with the highest frequency. In terms of indel, although the mutation frequency is not as high as SNV, generally one indel can produce about five or more neoantigen peptides.

**Table 2.**
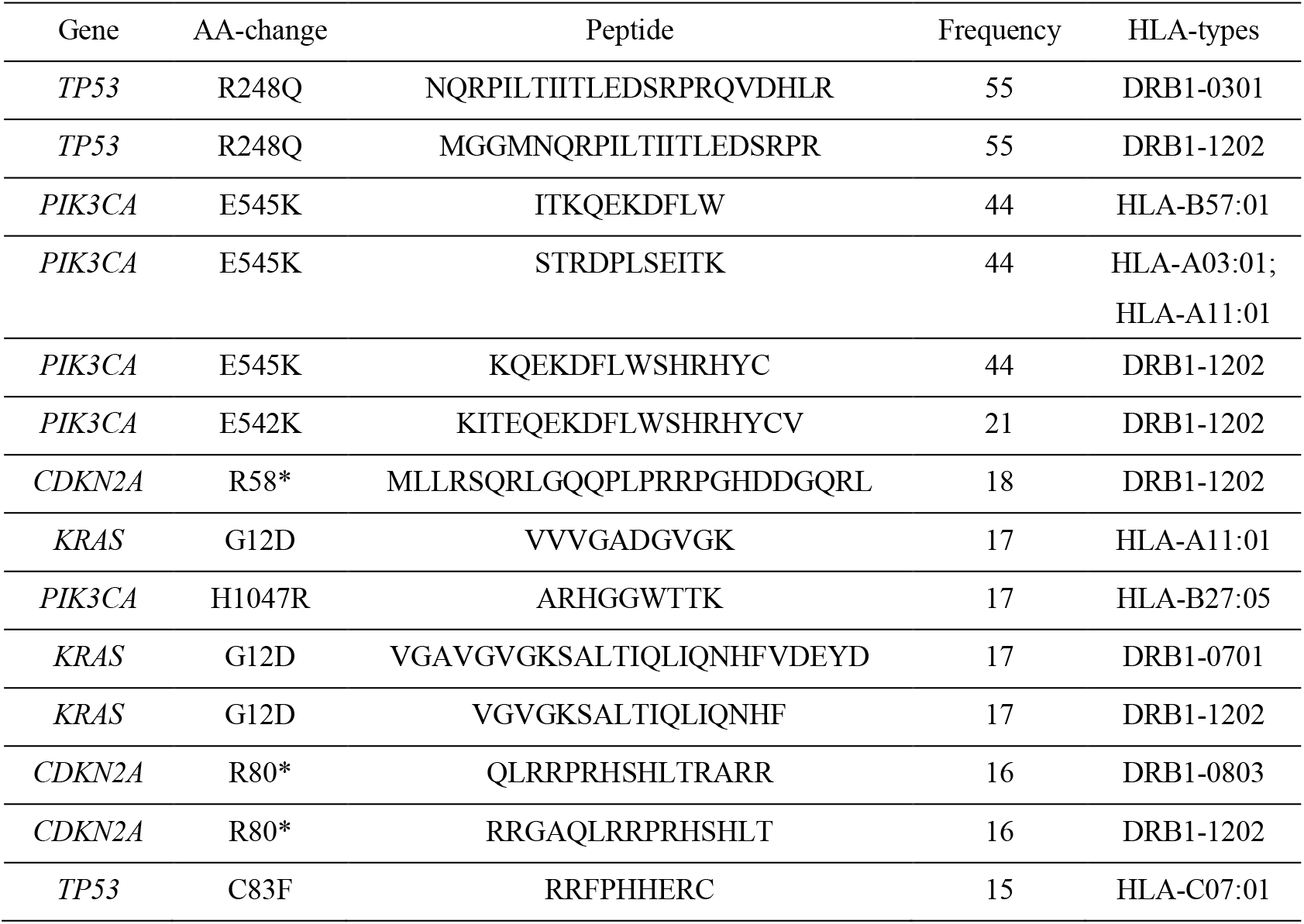
Top 15 SNV-related neoantigens in EC cohort.

**Table 3.**
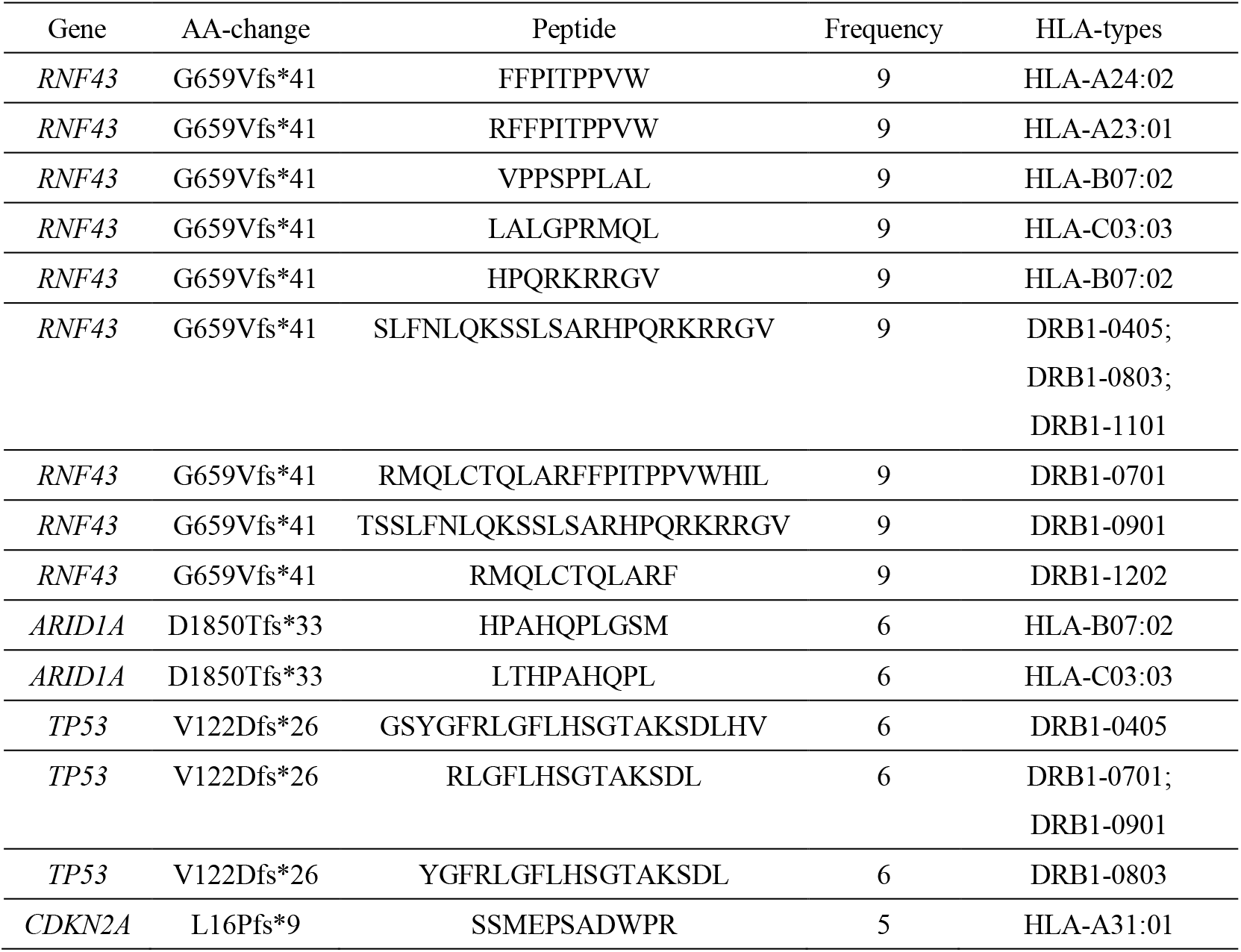
Top 15 INDEL-related neoantigens in EC cohort.

### 3.3. Comparison of Neoantigens in Different Subtypes and Cohorts of EC

By comparing the neoantigen profiles of different subtypes of EC, we found that, in the comparison of age classification, patients older than 60 had more neoantigens than younger patients, and men had more neoantigens than women, but they were not statistically significant (**Figure 2a-b**). In terms of cancer stage, Stage I EC carry the most abundant neoantigens, which may be related to the mutation load of the corresponding subgroup (**Figure 2c**). Notably, there were more neoantigens in ESCA than in ESCC patients (Fisher’s exact test, P<0.01, **Figure 2d**). Because the number of patients with known MSI status is small, we did not compare the neoantigen load of patients with different MSI status.

**Figure 2.**
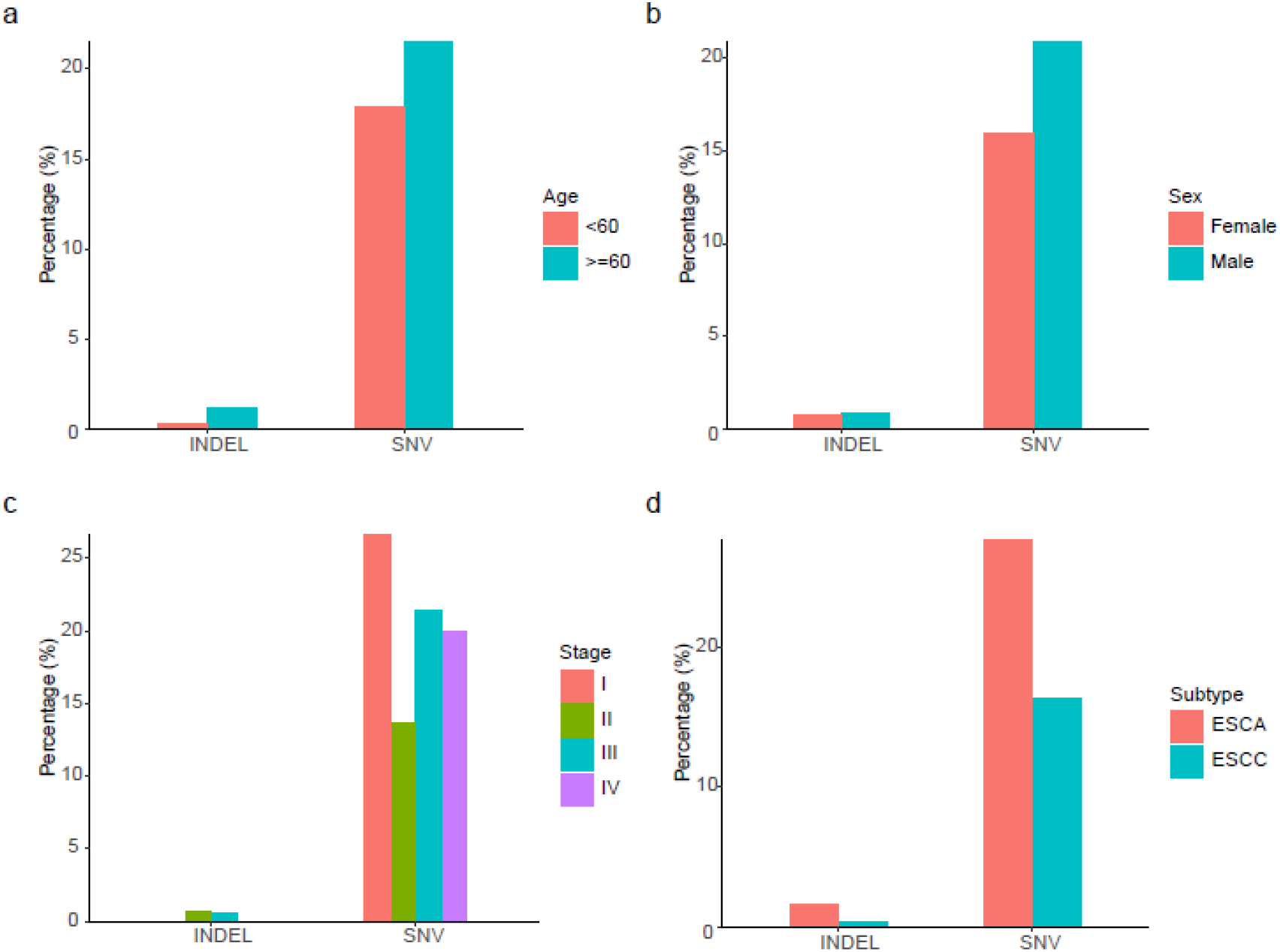
The comparation of neoantigens between different subgroups. (a) between age ≧ 60 and age <60 groups; (b) between female and male groups; (c) between different stage; (d) between ESCC and EAC subtypes. These analyses excluded patients with unknown subtypes.

**Figure 3.**
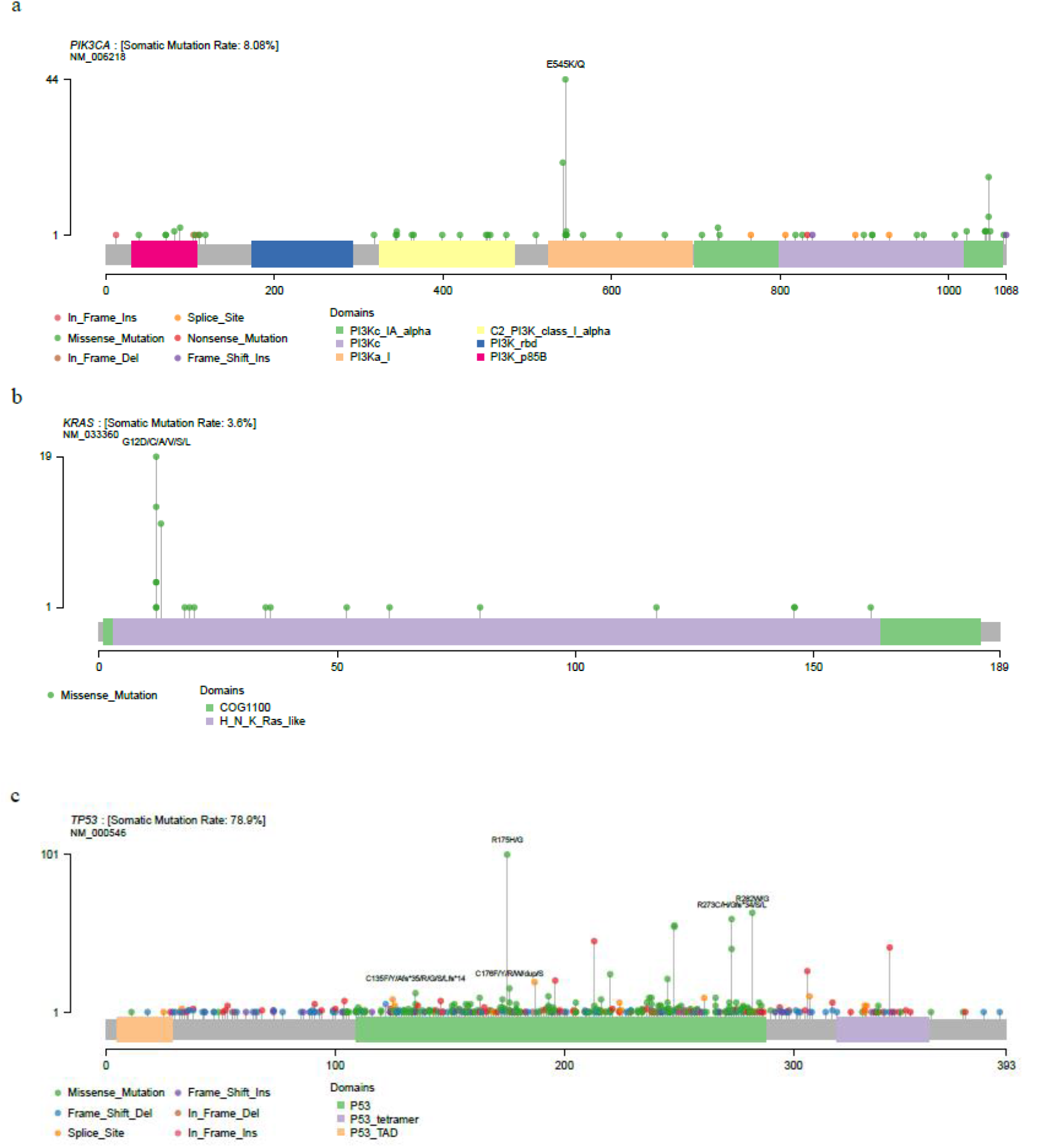
Mutational spectrum of *PIK3CA*(a), *KRAS* (b) and *TP53* (c) in 1886 EC patients.

### 3.4. Neoantigens Shared among EC Patients

To further investigate the potential significance of these high frequency neoantigens, we focused on sites with the highest mutation frequency, including R248Q (*TP53*), E545K (*PIK3CA*), G12D (*KRAS*), G12V (*KRAS*), and H1047R (*PIK3CA*), because these sites not only produce recurrent neoantigens, but also have a higher mutation frequency in EC cohort (Table 2).

*PIK3CA* is one of the hot driver genes of gastrointestinal malignancy [24, 27]. *PIK3CA* E545K is a hot spot mutation in several cancers, and two targeted drugs, Alpelisib and Fulvestrant, can target this mutation[28]. *PIK3CA* H1047R mutation is most commonly in breast cancer[29] and is also hot spots in multiple cancer types in the MSK-impact cohort[30]. Our previous studies have shown that this mutation is a potential neoantigen in patients with gastric cancer and colorectal cancer[14, 21]. Combined with the results of this study and previous studies, we suggest that this mutation may be a potential immunotherapeutic target for patients with gastrointestinal tumors.

*KRAS* Gly12 (including G12V, G12C, and G12D) is a classic cancer mutation in multiple cancers including endometrial, colorectal cancer, and non-small cell lung cancer[30]. We previously reported high frequency mutations and high frequency neoantigens of G12D/V in gastric and colorectal cancer[14, 31], our research group also demonstrated by mass spectrometry recently that *KRAS* G12V mutated neoantigen can be presented by HLA-A11:01 cell lines[32]. Previous studies have shown that gastrointestinal tumors, including esophageal cancer, gastric cancer, colon cancer and rectal cancer, have some similarities in tumor tissue origin and molecular characteristics[33], which are also reflected in this study, and high-frequency neoantigens in esophageal cancer are also similar to gastric cancer and colorectal cancer, suggesting the universality of neoantigen-based immunotherapy in gastrointestinal tumors.

In this study, we considered not only the class HLA-I alleles, but also the HLA-II alleles for potential neoantigens identification. The results showed that class HLA-II alleles (such as DRB1-0301, DRB1-1202) could also bind to lots of tumor-specific peptides, the number of which was comparable to that of the class HLA-I alleles. This indicates that HLA-II alleles should be considered as far as possible in the prediction of neoantigens in the future, so that we can expand our database of neoantigens and enable more patients to receive neoantigen-related immunotherapy.

## 5. Conclusions

Overall, based on 733 whole exome/genome sequencing data of EC samples and panel sequencing data, the most complete mutation landscape of EC was obtained. Based on the mutation and high-frequency HLA-I and HLA-II alleles, several recurrent neoantigens, such as neoantigens derived from mutations *PIK3CA* E545K, *KRAS* K12D, and *TP53* R248Q were identified. Some of these neoantigen-related mutations also have high frequencies in other gastrointestinal tumor, indicating that they are potential targets for gastrointestinal immunotherapy.

## Data Availability

Data supporting the results of this study are available from corresponding authors on request.

## Acknowledgments

We thank Ms. Xibao Song and professor Xiuqing Zhang for their support during this research.

