## Supplementary figures and images for "A Comprehensive Survey of Mutations in Oesophageal Carcinoma Reveals Recurrent Neoantigens as Potential Immunotherapy Targets"

### Supplemental figure S1

#### Mutation profiles of 733 WES esophageal carcinoma (EC) samples

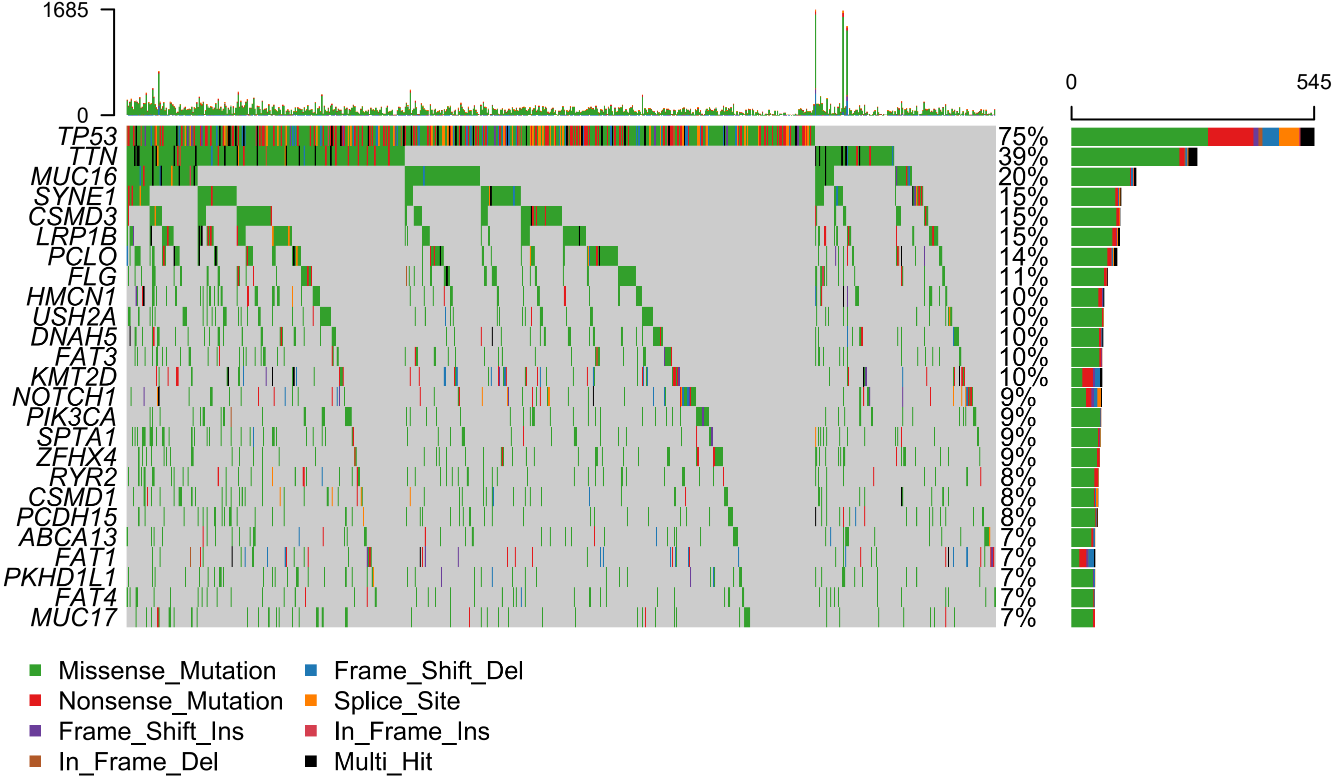

### Supplemental figure S2

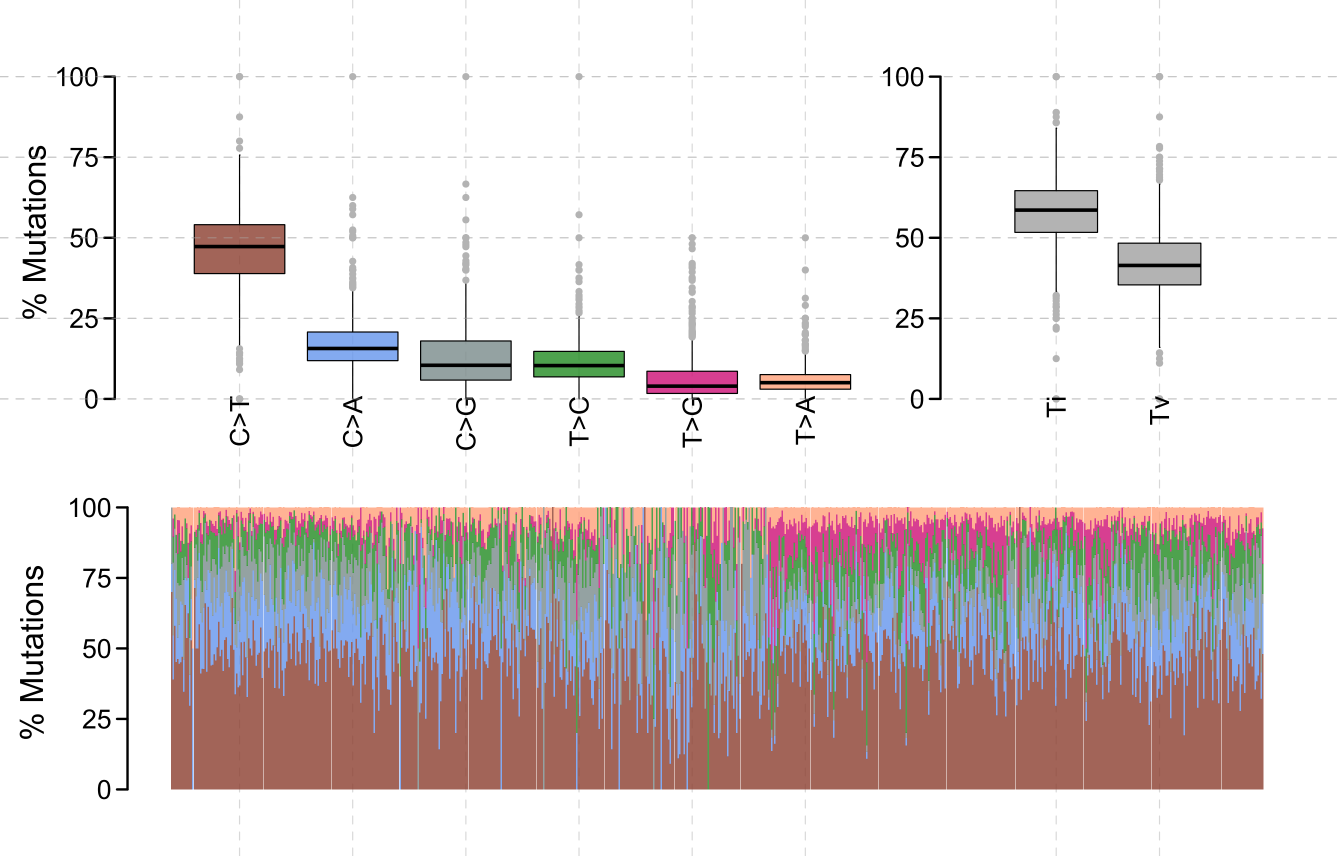

### Supplemental figure S3

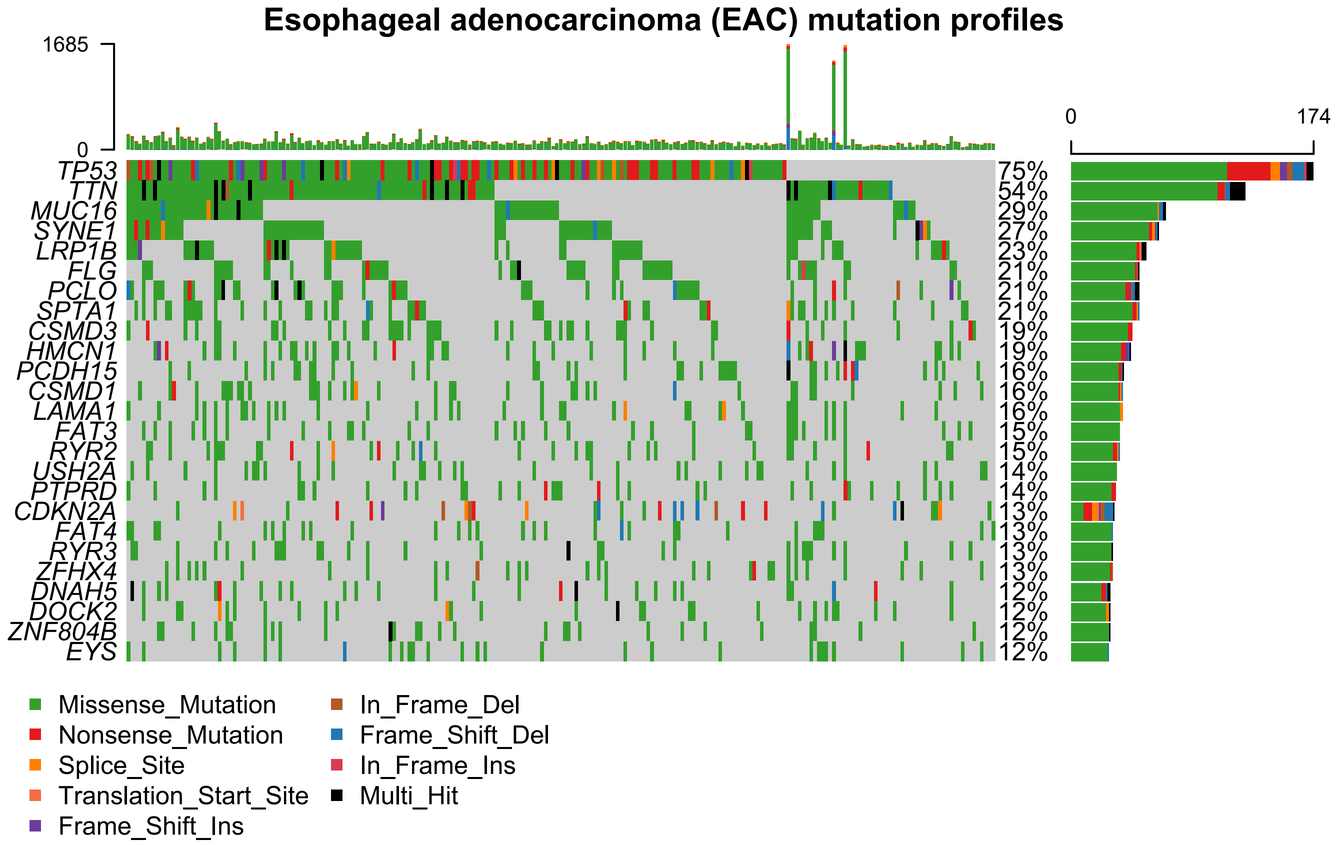

### Supplemental figure S4

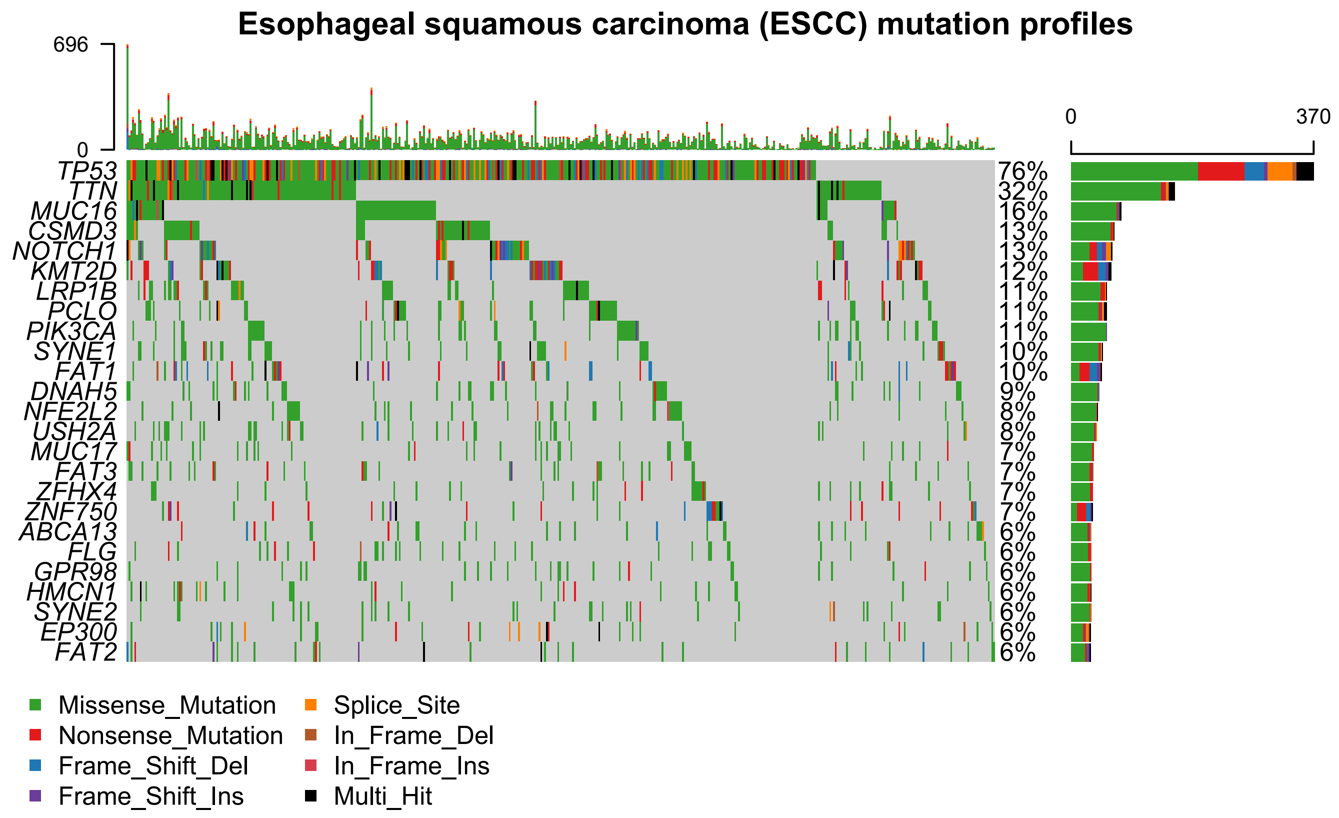

### Supplemental figure S5

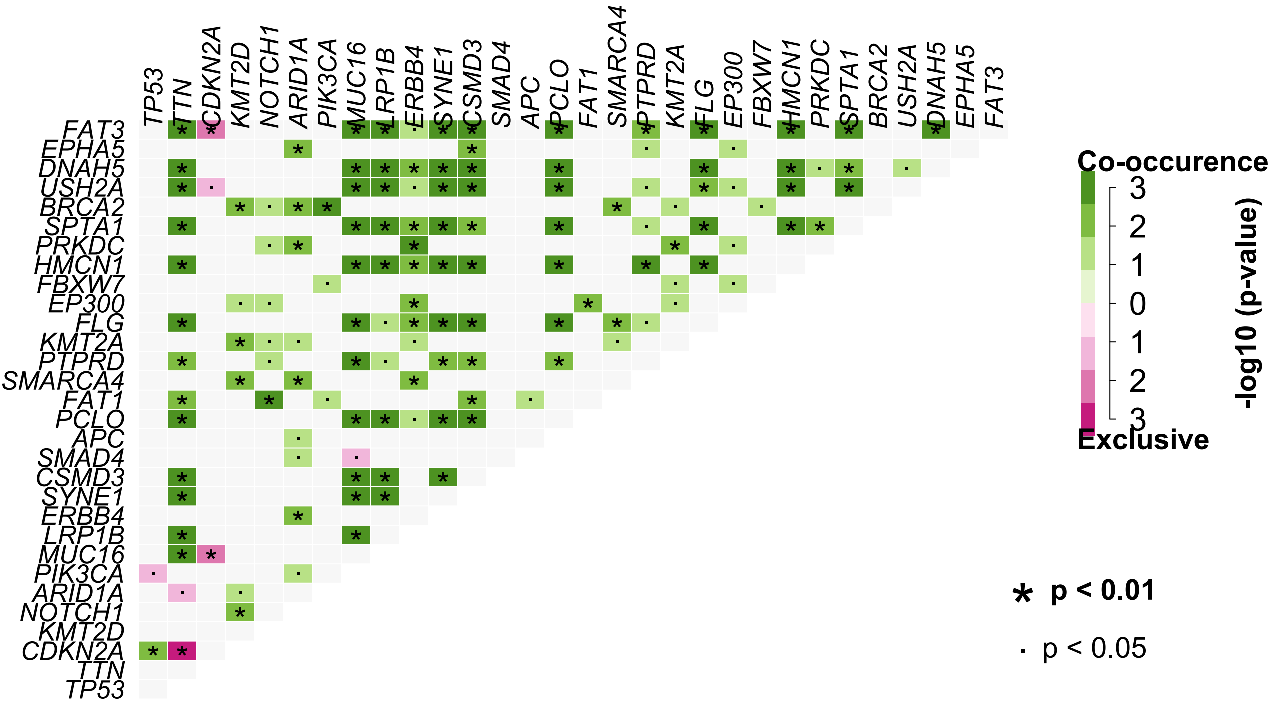
